# Zika virus outbreak in the Americas: Is *Aedes albopictus* an overlooked culprit?

**DOI:** 10.1101/044594

**Authors:** Azeem Mehmood Butt, Izza Nasrullah, Raheel Qamar, Yigang Tong

**Affiliations:** Genomics Laboratory, Department of Biosciences, COMSATS Institute of Information Technology, Islamabad 45550, Pakistan; Department of Biochemistry, Faculty of Biological Sciences, Quaid-i-Azam University, Islamabad 45320, Pakistan; State Key Laboratory of Pathogen and Biosecurity, Beijing Institute of Microbiology and Epidemiology, Beijing 100071, People’s Republic of China

**Author notes:** Co-first author. NOTE: An expanded version of this manuscript is under-preparation. The current pre-print version (v.01) of this manuscript may contain grammatical & proofreading mistakes. Errors and omissions excepted. Corresponding author: Azeem Mehmood Butt, PhD.

**Keywords:** Zika virus, codon usage, evolution, mutation pressure, natural selection

## Abstract

Codon usage patterns of viruses reflect a series of evolutionary changes that enable viruses to shape their survival rates and fitness toward the external environment and, most importantly, their hosts. In the present study, we employed multiple codon usage analysis indices to determine genotype specific codon usage patterns of Zika virus (ZIKV) strains from the current outbreak and those reported previously. Several genotype specific and common codon usage traits were noted in ZIKV coding sequences, indicative of independent evolutionary origins from a common ancestor. The overall influence of natural selection was found to be more profound than that of mutation pressure and acting on specific set of viral genes belonging to ZIKV strains of Asian genotype from the recent outbreak. Furthermore, an interplay of codon adaptation and deoptimization have been observed in ZIKV genomes. The collective findings of codon analysis in association with the geographical data of *Aedes* populations in the Americas suggests that ZIKV have evolved a dynamic set of codon usage patterns in order to maintain a successful replication and transmission chain within multiple hosts and vectors.

---

Zika virus (ZIKV), a member of the genus *Flavivirus* in the family *Flaviviridae,* is an enveloped, single-stranded positive-sense RNA virus. Like the closely related Dengue virus (DENV), ZIKA is also mosquito borne, spread by multiple *Aedes* species. Since its discovery in 1947, previous reports of ZIKV detection have been largely sporadic until the recent and ongoing outbreak of ZIKV in the Americas (Campos et al., 2015), with a few notable exceptions, including the ZIKV outbreaks in the Yap Islands (2007) (Duffy et al., 2009) and French Polynesia (2013–2014) (Cao-Lormeau and Musso, 2014). The close similarity between the clinical manifestations of ZIKV and those of DENV and Chikungunya virus (CHIKV) remains a potential obstacle to the efficient surveillance and management of ZIKV.

It is known that the genetic code shows redundancy and most of the amino acids can be translated by more than one codon. This redundancy represents a key step in modulating the efficiency and accuracy of protein production while maintaining the same amino acid sequence of the protein. Alternative codons within the same group that codes for the same amino acid are often termed ‘synonymous’ codons, although their corresponding tRNAs might differ in relative abundance in cells and in the speed by which they are recognized by the ribosome. However, the synonymous codons are not randomly chosen within and between genomes, which is referred to as codon usage bias (Grantham et al., 1980; Marin et al., 1989). This phenomenon has been observed in a wide range of organisms, from prokaryotes to eukaryotes and viruses. Studies on codon usage have identified several factors that could influence codon usage patterns, including mutation pressure, natural or translational selection, secondary protein structure, replication and selective transcription, hydrophobicity and hydrophilicity of the protein, and the external environment (Gu et al., 2004; Liu et al., 2011; Ma et al., 2013; Moratorio et al., 2013; Sharp et al., 1988; Tao et al., 2009). Moreover, considering the virus’s genome size and other viral features, such as dependence on host’s machinery for key processes, including replication, protein synthesis, and transmission, compared with prokaryotic and eukaryotic genomes, the interplay of codon usage among viruses and their hosts is expected to affect overall viral survival, fitness, evasion from host’s immune system, and evolution (Moratorio et al., 2013; Shackelton et al., 2006). Therefore, knowledge of codon usage in viruses can reveal information about molecular evolution as well as improve our understanding of the regulation of viral gene expression and aid in vaccine design, where efficient expression of viral proteins may be required to generate immunity.

In a recent *in silico* analysis, Freire *et al.* reported that the codon adaptation of the ZIKV non-structural 1 (NS1) gene is more biased towards *H. sapiens* than towards *Ae. aegypti* and suggested that this could improve viral replication within human cells (Freire et al., 2015). This is an interesting aspect of ZIKV evolution, however, the codon adaptation index analysis was only conducted in *H. sapiens* and *Ae. aegypti* and lacked some of the recently sequenced ZIKV genomes from the outbreak regions. As ZIKV is carried and transmitted by many types of *Aedes* mosquitoes other than *Ae. aegypti,* most notably *Ae. albopictus* (Duffy et al., 2009), understanding the evolutionary adaptation of viruses to their hosts and vectors is essential for developing effective surveillance, diagnostic, and preventive strategies. Based on the recently available sequence data from the outbreak and the broad range of the ZIKV vectors, we investigated the evolutionary adaptation of ZIKV to its transmission vectors and host to identify the potential cause of the ongoing massive ZIKV outbreak.

The workflow of phylogenetic and codon based analyses followed was similar as reported previously by us for CHIKV (Butt et al., 2014) and Marburg viruses (Nasrullah et al., 2015). The analyzed ZIKV genomes (n = 27) clustered into three separate groups of Asian, West African and East African genotypes where all of the recently reported outbreak sequences clustered into Asian genotype group. Although, we also noted that *NS1* gene had the highest adaptation towards *H. sapiens,* unlike the findings of Freire *et al.,* the high adaptation values of *NS1* were noted for all Asian isolates of pre‐ and present-endemic period. Moreover, *NS1* had similar codon adaptation index (CAI) values for *Ae. aegypti* and *H. sapiens.* Viruses such as ZIKV, DENV and CHIKV can replicate successfully in multiple hosts therefore, a virus may have to create a dynamic balance of both codon adaptation and codon deoptimization that could enable efficient replication and survival possibilities in multiple hosts of variable codon usage patterns. In such cases, we suspect that a single CAI observation is not enough to completely decipher the evolutionary patterns for such multi-host pathogen. We therefore, conducted similarity index and relative codon deoptimization index (RCDI) analyses to further investigate magnitude of ZIKV evolution under the influence of multiple hosts. ZIKV coding sequence were found to have highest codon deoptimization towards *Ae. albopictus* whereas, lowest codon deoptimization trends were noted for *Ae. aegypti* and *H. sapiens.* As reported previously, a low RCDI might indicate high adaptation to a host that comes in agreement to overall high CAI of ZIKV coding sequences towards *H. sapiens* and *Ae. aegypti.* On the other hand, a low RCDI is indicative that some viral genes are expressed in latency phases or even that the virus might present a low replication rate for successful establishment in a host with alternative codon usage patterns (Puigbo et al., 2010). Furthermore, according to the similarity index analysis, *Ae. albopictus* is probably the new preferred vector of ZIKV because the selection pressure exerted by *Ae. albopictus* on all three ZIKV genotypes is greater than the selection pressures imposed by *Ae. aegypti* or *H. sapiens* (Figure 1). The similarity index was found to be highest for *Ae. albopictus* whereas, lowest was noted for *H. sapiens*.

**Figure 1.**
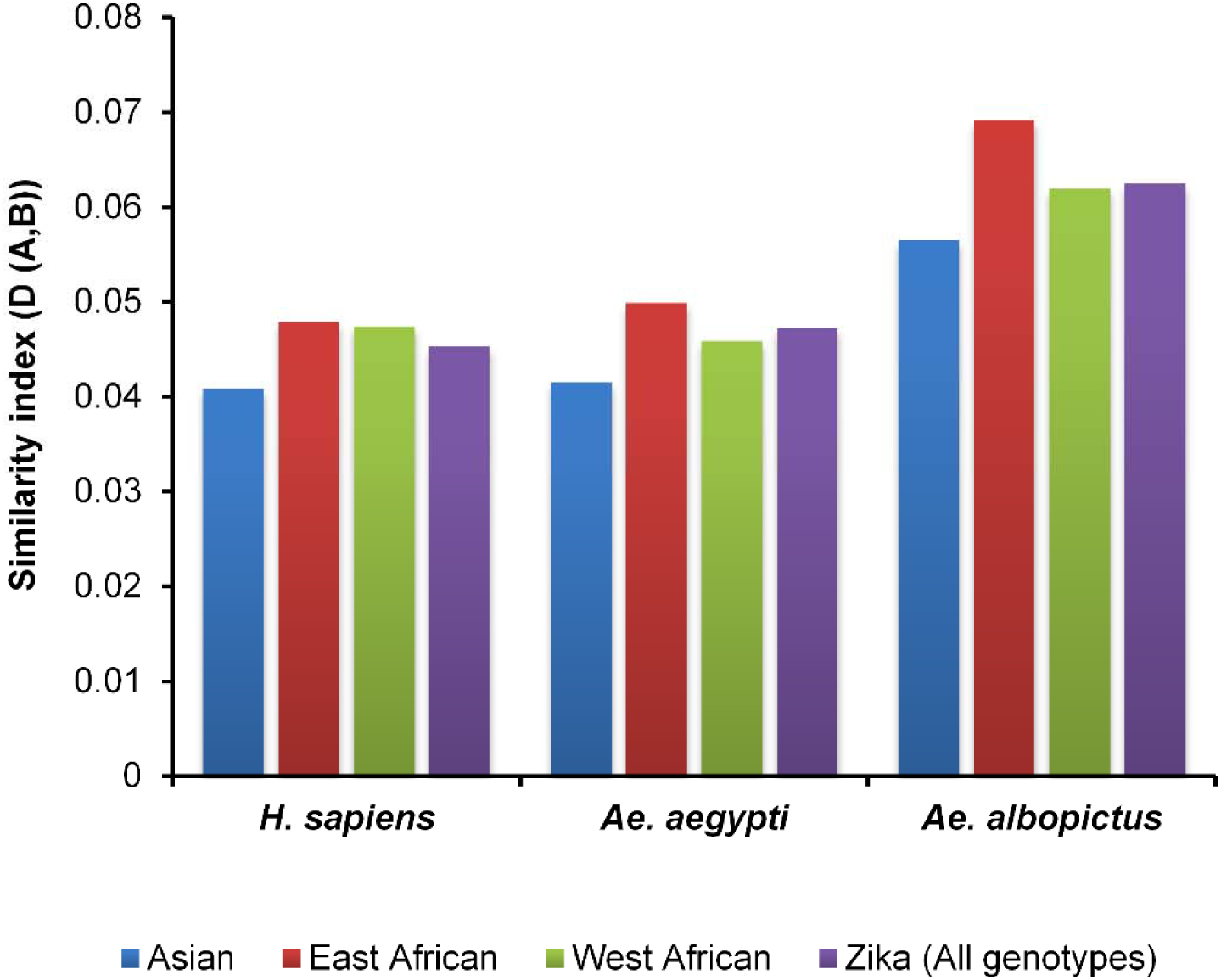
Analysis of host inducted selection pressure on coding sequences of ZIKV.

These observations are similar to that of adaptive evolution of CHIKV in *Ae. albopictus,* previously reported by us and others (Butt et al., 2014). Much convincing evidence has recently emerged to support this notion, including: (i) both ZIKV and CHIKV have been repeatedly observed to follow similar patterns of evolution and spread in different regions (Musso et al., 2015); (ii) in an environment containing both species of *Aedes* mosquitoes, a significantly greater number of *Ae. albopictus* than *Ae. aegypti* were ZIKV positive, and *Ae. albopictus* was identified as the primary vector for a ZIKV outbreak in Gabon (Grard et al., 2014); and (iii) a significant increase in the geographic distributions of *Ae. albopictus* in Asia, Africa, Europe, and the Americas was observed in a recent large-scale global distribution analysis (Kraemer et al., 2015). Presently, the major bottleneck that restricts complete understanding of ongoing outbreak is lack of availability of ZIKV full genome sequences. It is not known yet how ZIKV responds and manages within multiple vectors which are although member of same species but still demonstrates differences in certain aspects of genetic and environmental preferences. Currently, there is no reported genome sequence of ZIKV isolated from *Ae. aegypti* or *Ae. albopictus* populations of the affected areas. So, at this stage it is unclear if anyone or both types of *Aedes* are also actually harboring similar ZIKV strains as that of reported from the outbreak regions. By looking at previously reported ZIKV strains, there is only a single strain that was isolated from *Ae. aegypti* in 1966 (GenBank ID: HQ234499). Although, this strain clusters within Asian genotype group, by looking at correspondence analysis plots, it is clear that this strain contains several differences at individual gene levels from the rest of Asian strains (Figure 2). Whether these genome level differences are host specific warrants further investigations. In current scenario, host-vector-virus triangle highlights an important matter of concern towards reevaluating the current surveillance, eradication and preventative strategies for ZIKV which for now appear to be largely focused on *A. aegypti* only.

**Figure 2.**
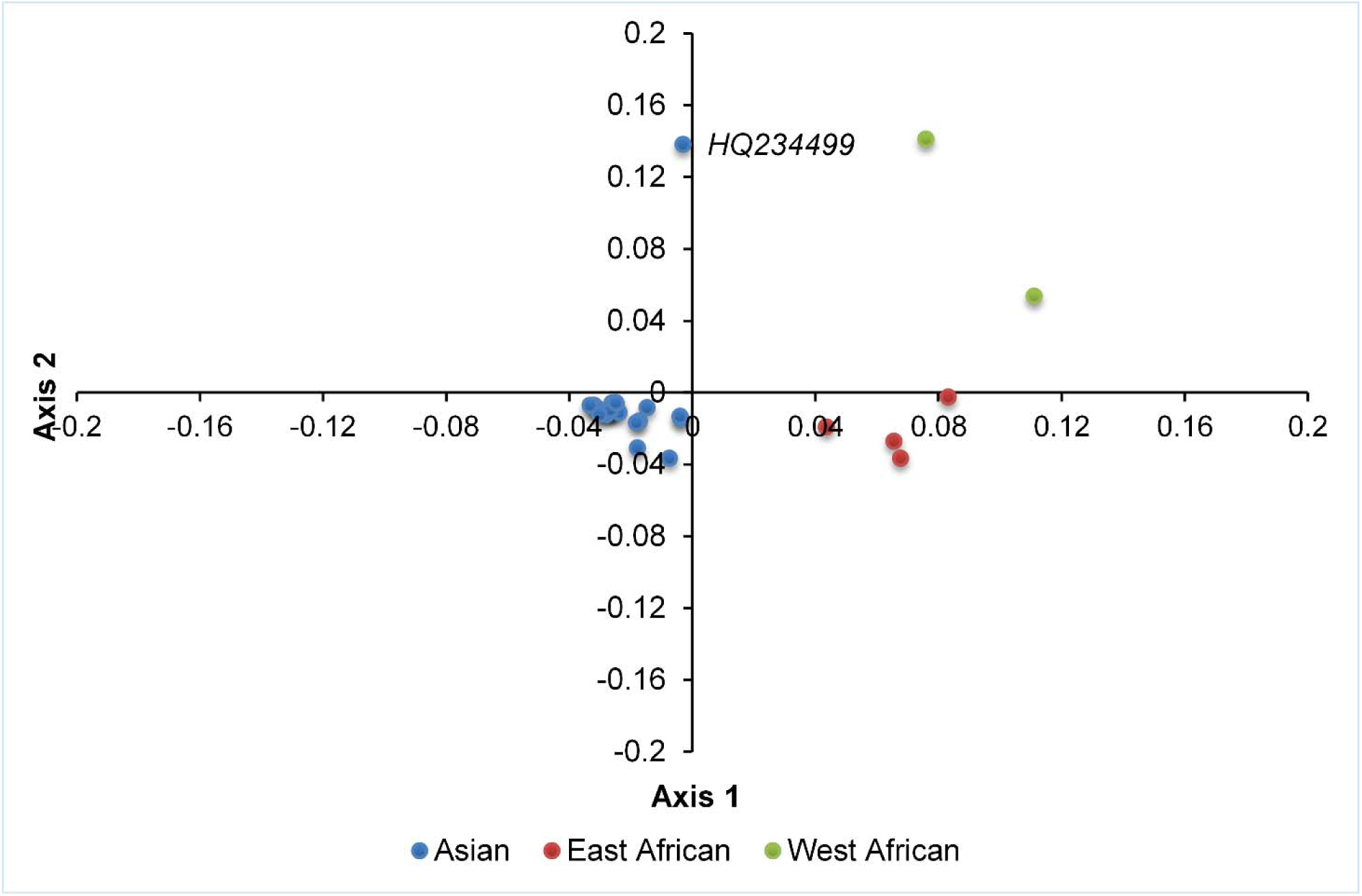
Correspondence analysis of codon usage pattern in ZIKV coding sequences. The single ZIKV strain from Ae. Aegypti is marked with its GenBank ID.

## Authors’ contributions

AMB RQ and YT conceived and designed the experiments. AMB and IN performed the experiments. AMB RQ and YT analyzed the data. RQ and YT contributed reagents/materials/analysis tools. AMB and IN wrote the paper. All authors read and approved the final manuscript.

## Acknowledgements

This research was supported by grant from China Mega-Project on Infectious Disease Prevention (No. 2013ZX10004-605).

